# City-wide metagenomics uncover antibiotic resistance reservoirs in urban beach and sewage waters

**DOI:** 10.1101/456517

**Authors:** Pablo Fresia, Verónica Antelo, Cecilia Salazar, Matías Giménez, Bruno D’Alessandro, Ebrahim Afshinnekoo, Christopher Mason, Gastón H Gonnet, Gregorio Iraola

**Affiliations:** Unidad de Bioinformática, Institut Pasteur Montevideo, Montevideo, Uruguay.; Proyecto “Centro de Metagenómica”, Institut Pasteur Montevideo, Montevideo, Uruguay.; Unidad de Microbiología Molecular, Instituto de Investigaciones Biológicas Clemente Estable, Montevideo, Uruguay.; Laboratorio de Calidad Ambiental, Intendencia Municipal de Montevideo, Montevideo, Uruguay.; Department of Physiology and Biophysics, Weill Cornell Medicine, New York, USA.; The HRH Prince Alwaleed Bin Talal Bin Abdelaziz Alsaud Institute for Computational Biomedicine, Weill Cornell Medicine, New York, USA.; The Feil Family Brain and Mind Research Institute, Weill Cornell Medicine, New York, USA.; ETH Zurich, Computer Science, Switzerland.; SIB Swiss Institute of Bioinformatics, Lausanne, Switzerland.; Centro de Biología Integrativa, Universidad Mayor, Santiago de Chile, Chile.

**Keywords:** Sewage, Beach, Metagenomics, Taxonomy, Antimicrobial resistance, Bacterial pathogens.

## Abstract

**Background:** Microbial communities present in environmental waters constitute a reservoir for antibiotic-resistant pathogens that impact human health. For this reason a diverse variety of water environments are being analyzed using metagenomics to uncover public health threats. However, the composition of these communities along the coastal environment of a whole city where sewage and beach waters are mixed, is poorly understood.

**Results:** We shotgun-sequenced 20 coastal areas from the city of Montevideo (capital of Uruguay) including beach and sewage water samples to characterize bacterial communities and their virulence and antibiotic resistance repertories. We found that sewage and beach environments presented significantly different bacterial communities. Sewage waters harbored a higher prevalence and a more diverse repertory of virulence and antibiotic resistant genes mainly from well-known enterobacteria, including carbapenemases and extended-spectrum betalactamases reported in hospital infections in Montevideo. Additionally, we were able to genotype the presence of both globally-disseminated pathogenic clones as well as emerging antibiotic-resistant bacteria in sewage waters.

**Conclusions:** Our study represents the first in using metagenomics to jointly analyze beaches and the sewage system from an entire city, allowing us to characterize antibiotic-resistant pathogens circulating in urban waters. The data generated in this initial study represent a baseline metagenomic exploration to guide future longitudinal (time-wise) studies, whose systematic implementation will provide useful epidemiological information to improve public health surveillance.

## Introduction

Human activity shapes the microbial communities residing in urban environments. In particular, urban sewage systems are designed to evacuate human wastes from the houses to areas of low human exposure and gradually reinstate them into natural watercourses such as creeks, beaches or the sea. This cycle is of tremendous importance for public health as waste waters can be reservoir and vehicle for the transmission of pathogenic bacteria and antibiotic resistance mechanisms. Indeed, the rapid emergence and spread of pathogenic bacteria with extensive antibiotic resistance has been recognized by the World Health Organization as a top health issue [1], since water can easily move microorganisms between humans and other animal species. Accordingly, the analysis of environmental waters is being adopted as an effective method to monitor the dynamics of antibiotic resistant pathogens [2], as this kind of environments can play a role as important as clinical settings for the selection of antibiotic resistance [3].

Recent advances in high-throughput sequencing (HTS) and computational biology now allow the exploration of microbial communities based on culture-independent approaches using metagenomics. This enables us to quantify and functionally characterize environmental microbiomes with unprecedented precision and comprehensiveness [4]. Indeed, the very recent implementation of this methodology to explore the microbial diversity in the built environment is providing a completely new layer of information to be integrated in the management of cities, potentially assisting decisions that range from urban design to public health [5,6]. In particular, urban sewage or beach water systems have been previously characterized using metagenomics not only focused on uncovering ecological patterns [7] but also in characterizing pathogenic and antibiotic resistant bacteria [8,9]. However, the joint analysis of bacterial communities present at the same time in the sewage and beach waters from the same metropolitan area remains to be explored in depth.

Sewage waters have been shown to accurately reflect the population’s gut microbiota composition [10], raising the possibility of using metagenomics to directly gain information about infection dynamics [11]. Additionally, beaches are important for recreational use but also are frequently recognized as risky environments for the contagion and transmission of bacterial infections [12], particularly if they are constantly or sporadically impacted by sewage spillovers. Accordingly, we performed a cross-sectional shotgun metagenomic analysis along the urban coast of Montevideo, the capital of Uruguay, aiming to characterize bacterial communities present in the sewage and beach water. Our study represents the first of that kind in a South American city and is the kick-off towards the incorporation of metagenomics in the surveillance of microbiological risks at city scale.

## Results

### Composition of sewage and beach communities

First, we explored the structure of microbial communities present in our beach and sewage samples using a multiset k-mer counting approach. This strategy provides an unbiased view that is not affected by taxonomic or functional assignment, conversely, it just evaluates the differential abundance of unique DNA segments [13]. Figure 1A shows a clustering analysis based on this methodology that shows a complete discrimination between sewage and beach samples, suggesting substantial differences in the composition of communities in these environments. Second, we confirmed the observed discrimination from a taxonomic point of view by calculating relative abundances of bacterial species present in each sample using an approach based on the identification and quantification of marker genes [14]. Figure 1B shows a dendrogram based on beta diversities (between samples) calculated using the Bray-Curtis dissimilarity distance from the taxonomic profiles, showing a complete discrimination between beach and sewage. Beta diversity was 0.42 within sewage samples (SD = 0.23) and 0.41 within beach samples (SD = 0.22) but increased to 0.63 (SD = 0.12) when comparing sewage against beach samples. Alpha diversity (within samples) was calculated using the Shannon index and averaged 3.65 (SD = 0.64) for sewage and 3.7 (SD = 0.4) for beach samples (Additional file 1: Fig. S1). These results indicate that taxonomic composition of bacterial communities from these environments are substantially different and can discriminate between beach or sewage origin (geographic location of each sample along the coast of Montevideo is displayed in Fig. 1C).

**Figure 1.**
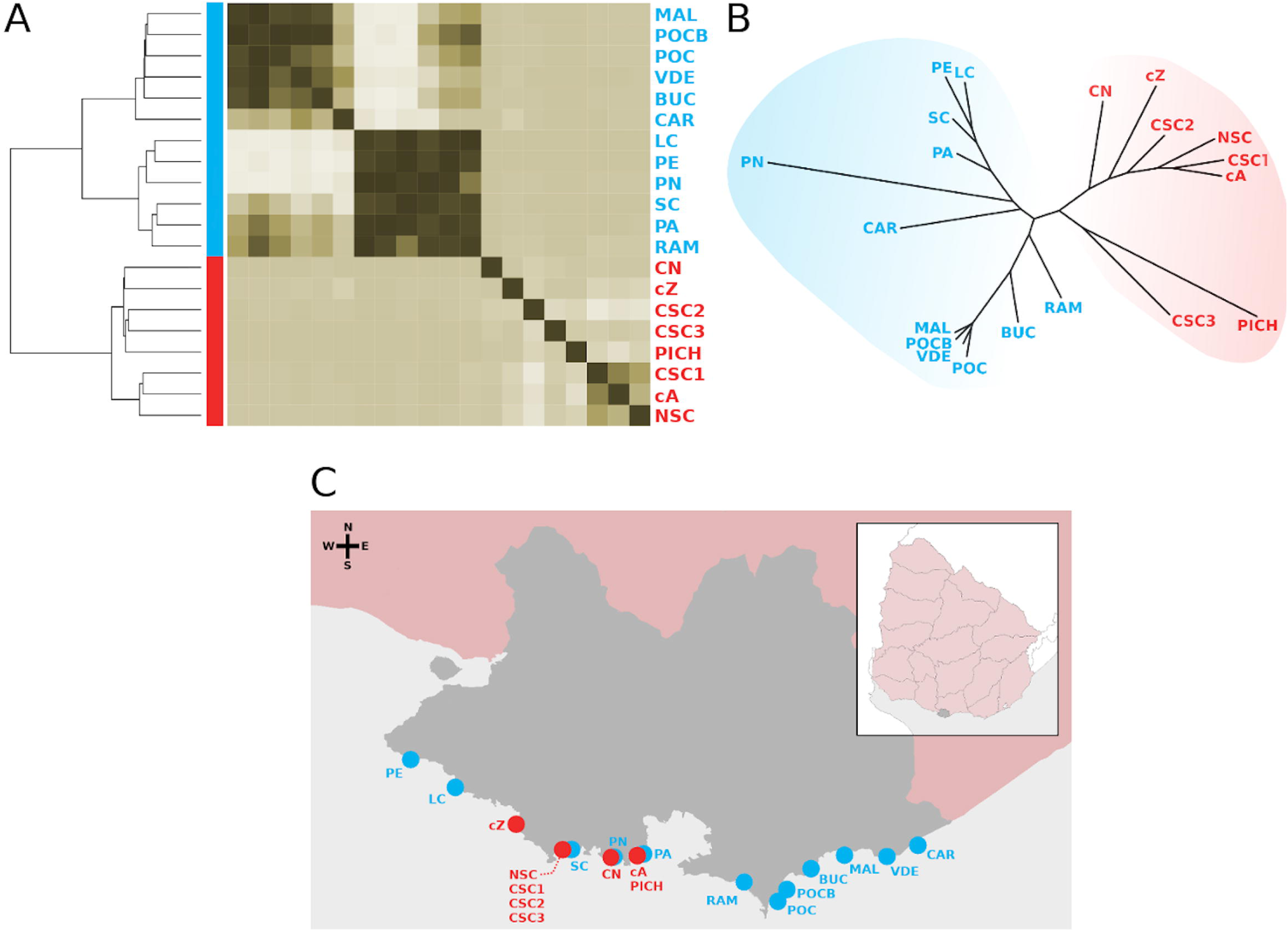
Community composition of beach and sewage waters of Montevideo. A) Heatmap showing a clustering analysis based on k-mer distances evidencing a complete separation between sewage (red) and beach (blue) samples. B) Clustering analysis separating sewage (red) from beach (blue) samples obtained by comparing beta diversities (dissimilarity between samples) calculated from relative abundance profiles of bacterial species. C) Sampling points along the coast of Montevideo (grey shade). Sewage water samples are in red and beach water samples are in blue.

### Occurrence of antibiotic resistance genes (ARGs)

The presence of different microbial communities in sewage and beach samples led us to hypothesize they could also encode distinct ARG repertories. Hence, metagenomic assemblies (Additional file 2: Table S1) were screened against the Comprehensive Antibiotic Resistance Database (CARD) [ 15] because is currently the most up-to-date and manually curated resource for ARGs detection. We found that 108 out of 2177 (~5%) ARGs had hits in our samples and they belong to 10 different antibiotic classes (Additional file 3: Table S2), being the clinically relevant TEM-4 and TEM-33 beta-lactamases the top occurring genes but aminoglycoside-modifying enzymes (like acetyl- or phosphotransferases) the most abundant class of ARGs. In particular, a significant difference (p = 0.002, Mann-Whitney U test) in ARGs abundance and a significantly higher diversity (p = 0.0024, Mann-Whitney U test) of ARGs according to the Simpson's index were found in favor of sewage compared to beach samples (Additional file 4: Fig. S2). Furthermore, when inspecting antibiotic classes we observed that sewage samples encoded ARGs belonging to 90% of antibiotic classes found in these environments while beach samples only encoded 40% of antibiotic classes, evidencing a more complex composition of antibiotic resistance mechanisms in the urban sewage waters (Fig. 2A). Indeed, only elfamycin resistance was present in beach but absent in sewage samples. On the other side, the occurrence of resistance mechanisms for aminoglycosides, betalactams, tetracyclines, sulfonamides, macrolides and streptogramins was significantly greater (p < 0.01, Mann-Whitney U test) in sewage compared to beach samples (Fig. 2B).

**Figure 2.**
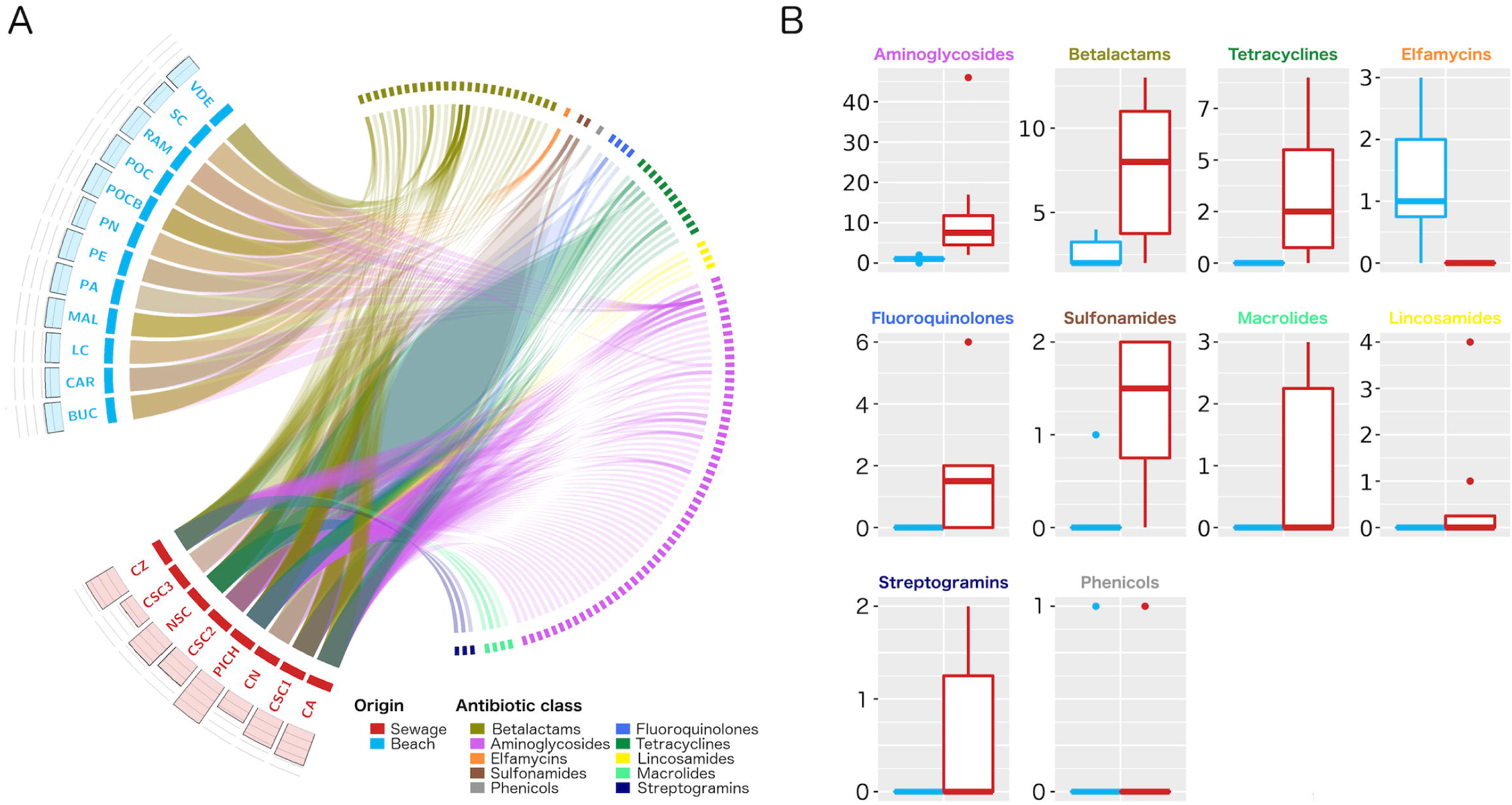
Occurrence of antibiotic resistance mechanisms. A) Circos representation showing the presence of ARGs across sewage (red) or beach (blue) samples. Links are drawn when a certain ARG (right blocks) is found in a certain sample (left blocks). Genes are colored according to antibiotic classes. Barplots above each left side block indicate the alpha diversity of ARGs within each sample. B) Boxplots showing ARG counts for different antibiotic classes in beach (blue) and sewage (red) samples.

### Distribution of ARGs in mobile elements

As a general trend, we found that ARGs present in our samples were more prevalent in plasmids than in bacterial chromosomes. Specifically, ARGs for sulfonamides, betalactams, aminoglycosides, phenicols, macrolides and streptogramins were more prevalent in plasmids than in bacterial chromosomes (Fig. 3A). Additionally, plasmids carrying ARGs found in our samples resulted to be extensively distributed among many clinically relevant enterobacteria like *E. coli*, *Salmonella*, *Klebsiella*, *Enterobacter*, *Citrobacter* and *Acinetobacter*, among others (Fig. 3B). ARGs for tetracyclines, lincosamides, fluoroquinolones and elfamycins were more frequently encoded in chromosomes.

**Figure 3.**
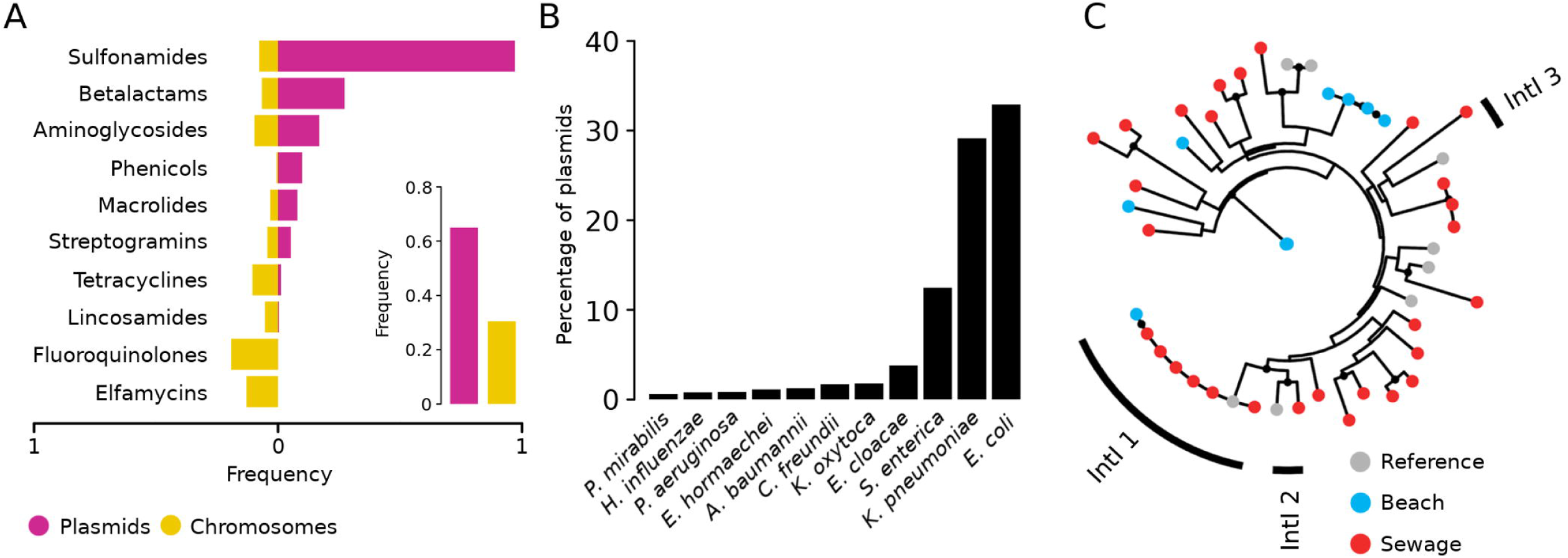
Distribution of ARGs in mobile genetic elements. A) Barplot showing the frequency of ARGs in bacterial chromosomes (yellow) or plasmids (violet) summarized by antibiotic class. B) Taxonomic distribution of bacterial plasmids carrying ARGs found in urban metagenomes. C) Phylogeny of reference integrase genes (grey) and those recovered from beach (blue) and sewage (red) samples.

We also looked for integrons and found a higher prevalence of them in sewage (74%) than in beach (24%) samples. Furthermore, clinically relevant integron classes 1, 2 and 3 were almost exclusively found in sewage samples (~90%) (Fig. 3C). Also, we were able to identify cassette ARGs with conserved *attC* sites associated to 24 out of 39 integrons (61%). These cassette genes mostly coded for multi-drug efflux pumps, but also we found carbapenemases (OXA family), GES extended-spectrum betalactamases (ESBLs), *aadA1* aminoglycoside nucleotidyltransferases, *cat* chloramphenicol acetyltransferases and *aac6-Ib* amikacin resistance. These cassette genes were exclusively found in sewage integrons.

### Occurrence of virulence factors (VFs)

To complement the characterization of ARGs we also screened metagenomic assemblies against the Virulence Factor Database (VFdb) [16]. Ninety nine out of 451 (~22%) VFs were detected in our samples. Specifically, VFs were found in 7 out of 8 (87.5%) sewage samples and 4 out of 12 (33%) beach samples. Sewage samples also presented a higher count and diversity of VFs compared to beach samples (Additional file 5: Fig. S3A). The functional classification of these VFs showed that those involved in bacterial motility, cell adherence, iron uptake and secretion were predominant among sewage samples. Interestingly, we found the presence of the ShET2 enterotoxin (*senB*) in a single sewage sample. We also examined the taxonomic distribution of these VFs and found that they are predominantly distributed in *Pseudomonas*, *Salmonella*, *Escherichia*, *Yersinia* and *Shigella*, among others (Additional file 5: Fig. S3B). These results highlight the importance of urban waters as reservoir and vehicle for VFs responsible of determining well-known pathogenic mechanism in clinically relevant bacteria.

### Taxa carrying mobile ARGs and VFs are sewage biomarkers

We also aimed to identify bacterial taxa associated to sewage or beach that can explain the observed differences in the composition of the overall bacterial community and their ARGs and VFs repertories. Then, we applied a linear discriminant analysis (LDA) and effect size estimation [17] to determine statistically significant taxa associated to beach or sewage. We summarized the results at the genus level and found 6 genera associated to the beach environment while 47 genera were characteristic of the sewage environment (Fig. 4A). Interestingly, 10 out of 47 (~21%) sewage-associated genera comprise bacterial species that are well-known human pathogens, including *Aeromonas*, *Acinetobacter*, *Arcobacter*, *Citrobacter*, *Enterobacter*, *Klebsiella*, *Pseudomonas*, *Sphingobacterium*, *Stenotrophomonas* and *Streptococcus* (Fig. 4B). Most of these taxa match with those found carrying mobile ARGs and VFs.

**Figure 4.**
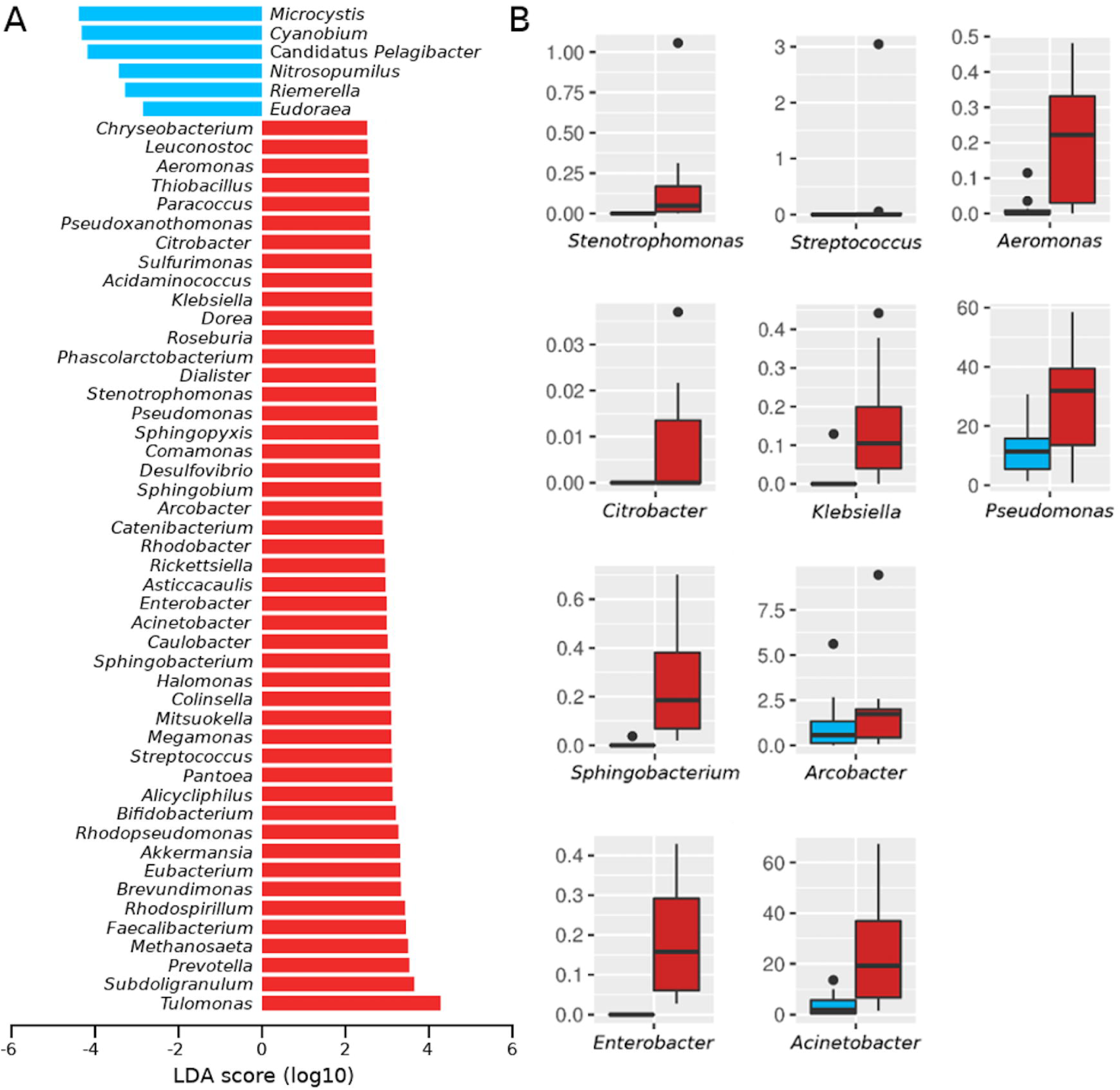
Identification of sewage biomarker taxa. A) Barplots showing LDA (Linear Discrimination Analysis) scores for bacteria genera that distinguish beach (blue) from sewage (red) samples. B) Boxplots comparing relative abundances between beach (blue) and sewage (red) samples for bacterial genera enclosing pathogenic species.

### Identification of pathogenic genotypes

To gain further insight on the microbiological risks in these environments, we attempted to resolve the presence of previously reported pathogens at the strain level using metagenome-derived multilocus sequence typing (MLST) [18,19]. Despite the fact that this method was unable to identify complete previously reported sequence types (STs), the detection of partial allele combinations allowed us to infer the presence of several genotypes of clinical importance. In sewage samples we found three alleles of *Citrobacter freundii* whose combination determines the ST-209, which has been detected from isolates recovered from diarrheal patients [20]. Also, we detected three alleles from *Escherichia coli* defining the clonal complex ST-131, which is a globally disseminated multidrug-resistant clone associated with human extraintestinal (urinary and bloodstream) infections [21]. Furthermore, the detection of two alleles of *Arcobacter cryaerophilus* defined the presence of ST-392, a recently characterized genotype causative of persistent diarrhea [22]. We also identified several *Salmonella* alleles that diverged from known genotypes, preventing the inference of putative STs. Many unknown alleles of *Pseudomonas fluorescens* were also identified both in sewage and beach samples. This species is mainly associated to food spoilage but has been sporadically reported as an opportunistic human pathogen causing systemic infections associated to the consumption of animal byproducts [23]. Overall, these results indicate the presence of pathogenic genotypes in urban waters of Montevideo.

## Discussion

Pharmaceutical products, including antibiotics, can be only partially metabolized by humans so these compounds or their derived metabolites are excreted [24] lastly reaching the environment. Sewage pipes have been largely considered as passive water transport systems, but recent studies uncovered hotspots of microbial diversity and activity in these urban environments. Indeed, the sewage system is the first step of city wastewater cycle and thus the most likely place where excreted antibiotic residues can induce the emergence of antibiotic resistant bacteria [25]. This is particularly relevant since recent studies have demonstrated that microbial communities present in the sewage recapitulate those found in the human gut microbiome [10]. Consequently, the exposure of human-derived bacteria to environmental pressures facilitates the emergence and spread of antibiotic resistant pathogens that can transmit and impact population’s health [26].

The characterization of urban waters in our capital city revealed a remarkably distinct taxonomic composition of bacterial communities found in sewage and beach environments, suggesting that the sewage system is efficient in evacuating most hazardous microorganisms far away from areas of human exposure. Indeed, the vast majority of virulence and antibiotic resistance mechanisms associated to clinically relevant pathogens were found in the sewage but not in beach samples. However, a more dense and longitudinal sampling is necessary to further characterize the dynamics of hazardous microorganisms circulating in these environments. Also, the comparison with metagenomes from hospital effluents along the city would provide a detailed picture of how nosocomial pathogens are being dispersed through the environment.

Indeed, many clinically relevant ARGs that we found in the city environment such as carbapenemases and ESBLs, have been frequently reported in nosocomial infections in Uruguay during the last decade [27–30]. This indicates that important antibiotic-resistant pathogens are being somehow transmitted among clinical settings and the urban environment, representing a public health threat. However, other ARGs such as metallo-β-lactamases that have also been reported in Uruguay [31–34] were not found in the sewage or beaches. Beyond technical biases are possible, this can be attributed to a differential capacity between distinct antibiotic-resistant clones to survive and spread in the environment; given that selective fitness of antibiotic-resistant pathogens (adapted to high antibiotic pressures in hospital settings) may be lower in less-exposed environmental waters [35].

In this sense, despite the urban environment may not be directly exposed to similar concentrations of antibiotics than those used to treat infections, sewage systems have been recognized as ARG reservoirs. So, considering that we found most environmental ARGs typically widespread in enterobacterial plasmids and that clinically relevant integrons were fundamentally recovered from sewage samples, genetic platforms for horizontal gene transfer can be playing a relevant role as reservoir of ARGs. Additionally, plasmids and integrons are prone to recombination [36] and genetic plasticity of certain bacteria has been proved to increase under subinhibitory pressures (as those probably found in the environment) with certain antibiotics [37], so the city sewage should be also considered as a birthplace for new antibiotic resistance mosaics mediated by recombination and horizontal gene transfer.

Regarding this, we were able to identify internationally disseminated pathogens like the *E. coli* ST-131 clonal complex, which encompasses multidrug-resistant genotypes with great capacity of recruiting new resistance genes. Also, we uncovered the presence of clinically underestimated bacteria like *Arcobacter cryaerophilus*, which is today considered an emerging waterborne pathogen and whose resistance to third generation cephalosporins has been already reported [38]. So, the compositional complexity of urban waters where different genotypes and gene repertories can coexist within a fluctuating bacterial community, opens the possibility of using environmental samples to monitor population’s health. Accordingly, beyond providing a detailed characterization of individual virulence and antibiotic resistance mechanisms, our study supports the possibility of using high-resolution metagenomics to study the epidemiological dynamics of antibiotic-resistant pathogens using urban waters as a proxy at the population-level.

## Conclusion

Our study represents a cross-sectional analysis of a metropolitan area encompassing more than 2.2 million inhabitants and, to the best of our knowledge, constitutes the first work using metagenomics to jointly characterize bacterial communities found in the sewage and beach waters of an entire city.

Our approach demonstrated its usefulness to identify antibiotic resistance determinants which were known to be present in nosocomial infections, as well as to uncover the presence of globally-widespread or underestimated pathogens with strain-level resolution. We consider that future longitudinal studies (time-wise) will be useful to monitor the fluctuations of bacterial communities, allowing the development of associative models with relevant metadata like outbreak information, rainfall or antibiotic prescription and stewardship.

The data generated in this initial study represent a baseline metagenomic characterization of environmental waters of Montevideo, which will be useful to guide future efforts to implement systematic studies aiming to evaluate antibiotic-resistant pathogen dynamics through time and space across different cities. This information can be later incorporated to improve public health surveillance for antibiotic-resistant pathogens.

## Methods

### Sample collection

We collected 20 water samples from 12 beaches and 8 sewage pipes or creeks where wastewater from the sewage system is poured directly. These sampling points are distributed along the whole coastal line of Montevideo, an uninterrupted system of sandy beaches that spans more than 20 kms (Fig. 1C, Additional file 2: Table S1). The samples were collected all the same day (around 3 hours elapsed from the first to the last). All samples were collected using sterile 200 mL plastic bottles and conserved in ice until they were processed within the same day.

### DNA purification and metagenomic sequencing

Each sample was centrifuged at 10,000 g for 15 min at 4 °C. Supernatants we discarded and pellets were processed using the FastDNA™ Spin Kit (MP Biomedicals) following the manufacturer's protocol. Paired end (2 × 125 bp) sequencing reads were generated on the Illumina HiSeq2500 machine, yielding in average 39.2 M (SD ± 5.4 M) reads per sample. On average 30.85 M (SD ± 4.07 M) reads passed an initial filter and were used in all further analyses. Sequencing data was deposited at the Sequence Read Archive (SRA) repository under BioProject number XXXXX.

### Metagenomic data analysis

Initial data quality inspection was performed with FastQC (https://www.bioinformatics.babraham.ac.uk/projects/fastqc) and then reads were filtered and trimmed using Trimmomatic [39] with the following parameters: LEADING:20, TRAILING:20, SLIDINGWINDOW:5:20, AVGQUAL:20 MINLEN:90. Resulting reads were used in downstream analyses. Characterization of bacterial pathogens at the strain-level was performed with metaMLST [19], that tries to identify multilocus sequence typing alleles directly from metagenomic sequences.

First, an unbiased description of the variability among communities in sewage and beach was obtained by running Simka [13] with default parameters. Second, MetaPhlan2 [14] was used to identify species and to determine their relative abundances across samples. Beta diversities were calculated over taxonomic profiles using the Bray-Curtis distance as implemented in the Vegan R package [40] and alpha diversities were calculated using the Shannon index in the base R package [41].

Metagenomes were *de novo* assembled for each sample with MEGAHIT [42]. Then, contigs over 1 kb. were retained and merged at 99% of identity using CD-HIT-EST [43]. Resulting contigs were secondary assembled using Minimus2 [44] requiring a minimum overlap of 100 bp. with at least 95% of identity at contig boundaries. Genes were predicted on the resulting contigs using Prodigal [45]. Antibiotic resistance and virulence genes were identified with Abricate (https://github.com/tseemann/abricate) by comparing contigs against CARD [15] and VFdb [16], respectively. Only hits with query coverage > 90% and sequence identity > 70% were kept.

Integrons were identified using IntegronFinder [46]. We used MAFFT [47] (with the L-INS-i option) to align the amino acid sequences of IntI genes recovered from metagenomes together with reference sequences of class 1 (IntI1, AAQ16665.1), class 2 (IntI2, AAT72891.1), class 3 (IntI3, AAO32355.1), class 4 (IntI4, AAC 38424) and class 5 (IntI5, AAD 55407.2) integrases. The resulting alignment was used to build a phylogenetic tree with RAxML [48].

## Declarations

### Ethics approval and consent to participate

Not applicable.

### Consent for publication

Not applicable.

### Availability of data and material

All data generated during this study is available at the Sequence Read Archive (SRA) under BioProject number XXXXX.

### Author’s contributions

GI and GG conceived the idea. BD, EA and CM generated the data. PF, VA, CS, MG and GI analyzed and interpreted the data. PF, VA, CS, GG and GI wrote the manuscript. All authors read and approved the final manuscript.

## Acknowledgements

We thank technicians from Intendencia de Montevideo for valuable assistance during sampling and data collection. We thank Hugo Naya for providing lab space and computational resources at the Bioinformatics Unit in the Institut Pasteur Montevideo during the development of this work.

